# Impact of Maternal Protein Restriction on Hypoxia-Inducible Factor (HIF) Expression in Male Fetal Kidney Development

**DOI:** 10.1101/2022.03.18.484938

**Authors:** Júlia Sevá Gomes, Letícia de Barros Sene, Gabriela Leme Lamana, Patricia Aline Boer, José Antônio Rocha Gontijo

**Author notes:** ***Ethics approval and consent to participate***: The Institutional Ethics Committee (CEUA/UNESP, Protocol #446) approved the experimental protocol; the general guidelines established by the Brazilian College of Animal Experimentation were followed throughout the investigation. ***Consent for publication:*** All authors approved for publication. **Data Availability:** Data are available in: http://repositorio.unicamp.br/acervo/detalhe/1166350?guid=1641498025205&returnUrl=%2fresultado%2flistar%3fguid%3d1641498025205%26quantidadePaginas%3d1%26codigoRegistro%3d1166350%231166350&i=1. ***Authors’ contributions:*** JSG: data curation, investigation, formal analysis, methodology, visualization, writing–original draft; GLL: methodology & visualization; LBS: methodology & supervision; PAB: conceptualization, formal analysis, methodology, supervision, visualization, writing–original draft & editing, JARG: conceptualization, funding acquisition, formal analysis, methodology, visualization, writing–review & editing. ***Data Availability Statement*** The miRNA sequencing data have been deposited into the Sequence Read Archive repository (accession: PRJNA694197) and Table S1 (Supplemental Information). ***Authors’ information***: José Antonio Rocha Gontijo, corresponding author, Department of Internal Medicine, School of Medicine, State University of Campinas, Campinas, SP, Brazil.

## Abstract

**Background:** The kidney ontogenesis is the most structurally affected by gestational protein restriction, reducing close to 30% of their functional units. The reduced nephron number is predictive of hypertension and cardiovascular dysfunctions generally observed in the adult age of most fetal programming models. We demonstrate HIF-1 predicts molecular pathway changes may be associated with the decreased nephron numbers in the 17 gestational days (17GD) low protein (LP) intake male fetal kidney compared to regular protein (NP) progeny intake. Here, we evaluated the predicted targets and related pathways in the fetal kidneys of 17-GD LP offspring to elucidate the molecular modulations during nephrogenesis.

**Methods:** Pregnant Wistar rats were allocated into two groups: NP (regular protein diet −17%) or LP (diet-6%). Taking into account miRNA transcriptome sequencing previous study (miRNA-Seq) in 17-GD male offspring kidneys using previously described methods, was investigated predicted target genes and proteins, in particular, related to HIF-1 pathway by RT-qPCR and immunohistochemistry.

**Results and conclusions:** The current study data supported that nephron onset impairment in the 17-DG fetus’s kidney, programmed by gestational low-protein intake, is, at least in part, related to alterations in the HIF-1α signaling pathway. Factors that facilitate the transposition of HIF-1α to the mesenchymal cell’s nucleus, such as NOS, Ep300, and HSP90, may have an essential role in this regulatory process. This alteration leads to the inhibition of adaptive responses to the adverse environment, secondary to an increase in ungraded HIF-1α, possibly associated with a reduction in the transcription factor elF-4 and proteins of their respective signaling pathways. Consequently, we may suggest an early maturation process of renal cells, inhibition of nephron progenitor cell division, and reduction of renal functional units in the offspring of rats submitted to severe gestational protein restriction.

## INTRODUCTION

Embryo/Fetal programming is caused by psychological, placental ischemia, and nutritional stress during development, leading to long-term effects on different organ structure and function disorders with an increasing predictive chance of developing the chronic disease [1–9]. Thus, it may affirm that the intrauterine environment regulates fetal growth trajectory and predicts future diseases. The gestational nutritional restriction results in several changes in the fetal organs and systems during developmental stages, which may cause disorders in adult life [5–8, 10–12]. Previous studies have demonstrated that fetal programming results in low birth weight, fewer nephrons, and increased risk of cardiovascular and renal disorders in adulthood [3–8, 13]. The low nephron number is related to arterial hypertension, and in hypertensive patients, approximately 40% of the number of nephrons is reduced [8,14]. Prior experimental studies from our lab and other authors have demonstrated lower birth weight, 28% fewer nephrons, reduced renal salt excretion, chronic renal failure, and enhanced systolic pressure from 8 to 16 weeks of life in gestational low-protein (LP) intake compared to standard (NP) protein intake offspring in adulthood [3–7, 15]. However, information regarding the molecular mechanisms of the etiopathogenesis of nephrogenesis cessation is still scarce. Nephrogenesis involves fine control of gene expression, protein synthesis, tissue remodeling, and cell fates of the different kidney progenitor cells [16] Huang et al., 2020]. During renal ontogenesis, nephron stem cell renewal and differentiation are too controlled to generate an adequate number of nephrons. The kidney nephron numbers are defined by a closed interaction among ureter bud (UB) and metanephric mesenchyme (MM) progenitor cells [8, 17–19]. Signals from MM induce UB-stimulated growth and branching of the tubule system. In turn, MM proliferation and differentiation, constituting a mesenchymal cap (CM), are mediated by UB ends [20]. There has been serious interest in the role of epigenetic impact, concerning the long-term effects of prenatal stress, on fetal development. MicroRNAs (miRNAs) are genome encoded small non-coding RNAs of approximately 22 nucleotides in length and play an essential role in the post-transcriptional regulation of target gene expression [21–23]. Studies indicate that miRNAs are involved in many regulatory biological networks during development and cell physiology. Thus, miRNAs characterization is indispensable during nephron ontogenesis and may help us understand gene regulation and cellular proliferation, differentiation, and apoptosis and explain the pathophysiology, including kidney disorders [24–29] in adulthood. We recently demonstrated in the fetus 17 days of gestation (GD) protein-restricted male fetus changes in metanephros miRNAs. We predicted mRNA expression that encodes proteins related to a 28% reduction in nephrogenic stem cells in the cap metanephric (CM). Thus, could be suggested that miRNAs, mRNAs, and protein disruption could have reduced proliferation and promoted early cell differentiation [7]. Hypoxia-Induced Factors (HIF) are transcriptional factors from the helixloop-helix-PAS family consisting of labile α unit, and stable beta unit to form a transcriptional complex. The heterodimer translocate to the cell nucleus activating several target genes [30,31]. For many ways is the HIF activity modulated. At normal tissue oxygen level, HIF-1α is hydroxylated and recognized by the VHL E3 compound (tumor suppressor Von Hippel Lindau, Culina 2, Elonguin C, and Rbx1), which promotes its degradation in the proteasome. Otherwise, in low oxygen tissue tension or the absence of VHL protein, HIF-1α escapes degradation [30, 31]. On the other hand, the activation of HIF-1α occurs in connection with tumor suppressor p53 mediated by ubiquitin MDM-2, also leading to degradation in the proteasome. Consequently, decreases expression of p53 also leads to reduce HIF-1 degradation. The main p53 function is the maintenance of the integrity of the genetic code whose structure must be constant in different cells of the organism by a set of reactions that activate repair proteins or block gene changes. In case of DNA extreme damage, the p53 protein prevents cell mitosis and completes cell division achieving death through apoptosis or the impediment of these cells to multiply definitively, causing cellular senescence [32,33]. HIF-1α is also degraded by the action of the chaperone protein HSP-90, through its conformational alteration.–The signaling pathway of PI3K/AKT and mTOR was also demonstrated as activating the expression of HIF-1α, both, in an oxygen-rich and hypoxia environment while HIF is activating by the elF-4 factor in normal oxygen levels [34,35]. Several genes are targets for the HIF-1α as a transcriptional factor such as erythropoietin and transferrin expression, activation of α and beta transformer growth factor (TGFα and TGFβ), as well the activation of endothelial vascular growth factor (VEGF) and nitric oxide synthetize (NOS2). During embryonic development, HIF-1α is identified in virtually all tissues, and specifically in the kidney, was identified in the distal tubules and collecting ducts and in a smaller amount in the peripheral cortex. In the HIF-1α knockout rats no response of VEGF production during hypoxia, compromising angiogenesis and leading to the death of the animal in the period of development demonstrating the importance that HIF-1α on renal development. However, there is still no consolidated study, in animal programming models whose mother was submitted to gestational protein restriction, demonstrating the implication of the HIF-1α signaling pathway on the nephrogenesis process. So, in the current study, we evaluated, in the last gestational days, the miRNAs transcriptome and predicted targets of those to try to elucidate the molecular implication on nephrogenesis in male offspring submitted to gestational low protein diet.

## MATERIAL AND METHODS

### Animal and Diets

The experiments were conducted as described in detail previously [Mesquita et al., 2010a,b] on age-matched female and male rats of sibling-mated Wistar *HanUnib* rats (250–300 g) originated from a breeding stock supplied by CEMIB/ UNICAMP, Campinas, SP, Brazil. The environment and housing presented the right conditions for managing their health and well-being during the experimental procedure. Immediately after weaning at three weeks of age, animals were maintained under controlled temperature (25°C) and lighting conditions (07:00–19:00h) with free access to tap water and standard laboratory rodent chow (Purina Nuvital, Curitiba, PR, Brazil: Na+ content: 135 ± 3μEq/g; K+ content: 293 ± 5μEq/g), for 12 weeks before breeding. The Institutional Ethics Committee on Animal Use at São Paulo State University (#446-CEUA/UNESP) approved the experimental protocol, and the general guidelines established by the Brazilian College of Animal Experimentation were followed throughout the investigation. It was designated day 1 of pregnancy as the day in which the vaginal smear exhibited sperm. Then, dams were maintained ad libitum throughout the entire pregnancy on an isocaloric rodent laboratory chow with either standard protein content [NP, n = 36] (17% protein) or low protein content [LP, n = 51] (6% protein). The NP and LP maternal food consumption were determined daily (subsequently normalized for body weight), and the bodyweight of dams was recorded weekly in both groups. On 17 days of gestation (17-DG), the dams were anesthetized by ketamine (75mg/kg) and xylazine (10mg/kg), and the uterus was exposed. The fetuses were removed and immediately euthanized by decapitation. The fetuses were weighed and, the tail and limbs were collected for sexing. The metanephros was collected for Next Generation Sequencing (NGS), RT-qPCR, and immunohistochemistry analyses.

### Sexing determination

The present study was performed only in male 17-GD progeny, and the sexing was determined by Sry conventional PCR (Polymerase Chain Reaction) sequence analysis. The DNA was extracted by enzymatic lysis with proteinase K and Phenol-Chloroform. For reaction, the Master Mix Colorless—Promega was used, with the manufacturer’s cycling conditions. The Integrated DNA Technologies (IDT) synthesized the primer following sequences below:

1. Forward: 5’-TACAGCCTGAGGACATATTA-3’
2. Reverse: 5’-GCACTTTAACCCTTCGATTAG-3’.

### Total RNA Extraction

RNA was extracted from an isolated two-kidney pool from each litter of the NP (n = 5) and LP (n = 5) offspring using Trizol reagent (Invitrogen), according to the instructions specified by the manufacturer. After centrifugation, the material is separated into 3 phases, (a) the upper phase is aqueous and clear; (b) the intermediate phase, and (c) the reddish lower organic phase. The RNA remains in the aqueous phase and, being recovered through precipitation, carried out by washing isopropyl alcohol cycles, and centrifugation. Total RNA quantity was determined by the absorbance at 260 nm using nanoVue spectrophotometer (GE Healthcare, USA), and the RNA purity was assessed by the A 260 nm/A 280 nm and A 260 nm/A 230 nm ratios (acceptable when both ratios were >1.8). RNA Integrity was ensured by obtaining an RNA Integrity Number - RIN >8 with Agilent 2100 Bioanalyzer (Agilent Technologies, Germany).

### Real-time Quantitative PCR (mRNAs)

For the analysis of expression level of NOS2, p53, HSP90, HIF-1α, NFκB, elF4, Ep300, TGFβ-1, mTOR, AT1α, AT1β, and AT2, in the isolated two-kidney pool, RT-qPCR was carried out with SYBR Green Master Mix, using primers specific for each gene, provided by Exxtend (Campinas, SP, Brazil) (Table 1). Reactions were set up in a total volume of 20 μL using 5 μl of cDNA (diluted 1:100), 10 μL SYBR Green Master Mix (Life Technologies, USA), and 2.5 μL of each specific primer (5 nM) and performed in the StepOnePlusTM Real-Time PCR System (Applied BiosystemsTM, USA). The cycling conditions were 95°C for 10 minutes; 45 cycles of 95°C for 15 seconds and 60°C for 1 minute. Ct values were converted to relative expression values using the ΔΔCt method with offspring kidneys data normalized to GAPDH as a reference gene.

**Table 1.**
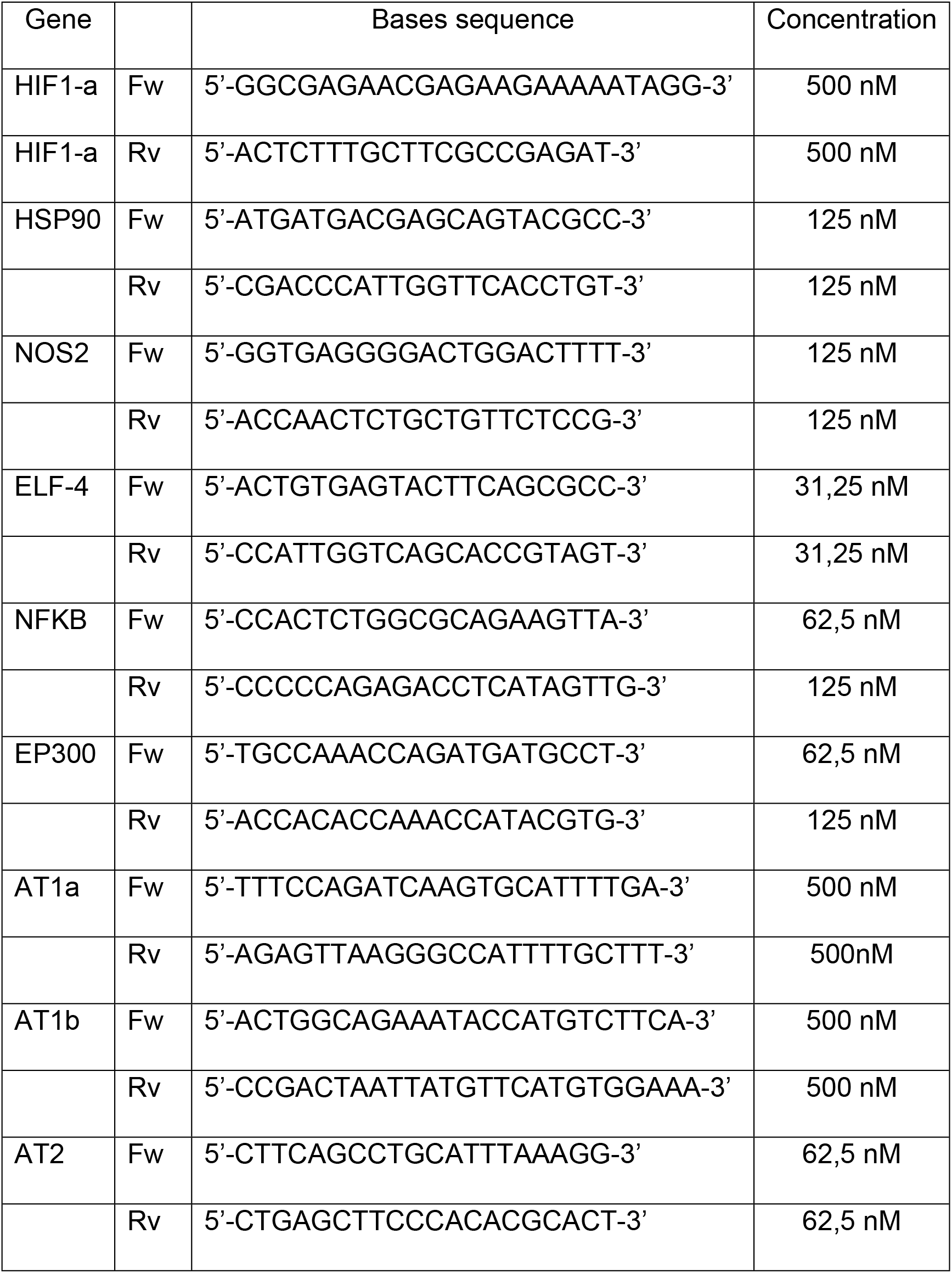
The sequence of the primers used for RT-qPCR was designed by the company IDT.

### Analysis of the Gene Expression

To analyze the differential expressions, the miRNA or mRNA levels obtained for each gene (Table 1) were compared with the LP group concerning the appropriated NP group. Normalization of miRNA expression was made using the expression of the snRNA U6 and snRNA U87 reference genes (Accession: NR_004394 and AF272707, respectively), and, for mRNA expression, the genes GAPDH, β-actin, and TBP. Relative gene expression was evaluated using the comparative quantification method. All relative quantifications were assessed using DataAssist software v 3.0, using the ΔΔCT method. PCR efficiencies calculated by linear regression from fluorescence increase in the exponential phase in the program LinRegPCR v 11.1 [5,7].

### Immunohistochemistry

After perfusion, the kidneys from NP (17-GD, n=5 from different mothers), and LP (17GD, n=5 from different mothers) offspring were removed and placed in the same fixative for 2 h, followed by 70% alcohol until processed for paraffin inclusion or frozen in liquid nitrogen. The frozen and paraffin blocks were cut into 5-μm-thick sections. For immunofluorescence, the paraffin sections were incubated sequentially with: (1) phosphate-buffered saline (PBS) containing 5% milk for 45 min to minimize non-specific reactions, (2) incubated overnight at 4°C, with primary antibodies for anti-HIF-1α, anti-HSP-90, anti-NOS2, anti-elF-4, anti-p-elF4, anti-TGFβ-1, anti-mTOR and anti-NFκB (Table 2) diluted in 1% BSA, and (3) goat anti-mouse Alexa 488 antibody (Alexa) for 2 h at room temperature. Secondary antibodies were used according to the primary antibody. Finally, sections were revealed with DAB (3,3’-diaminobenzidine tetrahydrochloride, Sigma - Aldrich CO^®^, USA), counterstained with Mayer’s hematoxylin, dehydrated, and mounted. After incubation, the sections were rinsed in 0.1 M PBS and cover-slipped with Vectashield anti-fading medium containing DAPI (Vector). No immunoreactivity was seen in control experiments in which one of the primary antibodies was omitted. The images were obtained using the photomicroscope (Olympus BX51). *Morphology quantification* - Hematoxylin-eosin stained paraffin sections (NP and LP from different mothers, [n=5]) were used to measure renal, cortical, and medullar areas in the kidneys from 17-GD offspring. In paraffin 5μm kidney histological sections in HE, the CAP areas and cell analysis were performed by microscopic fields digitized (Olympus BX51) using CellSens Dimension or *ImageJ* software. The relative percentages of the cortical and medullary area relative to the total renal area, previously determined, were also determined. The whole kidney area corresponds to the determination of measures of the entire cut, that is, the sum of cortical and medullary areas. Therefore, we reiterate that the histological section’s whole CAP regions were evaluated for each animal (n = 5) in both groups. For this procedure, the kidneys were sectioned longitudinally in half and embedded in paraffin each half. Then, the microtome cuts were made after the block was trimmed and stained in HE. The studies were carried out blindly and similarly for both groups of animals (NP and LP).

**Table 2.** Dilution of antibodies used in immunohistochemistry.

| Primary antibody | Dilution | Antibody supplies |
| --- | --- | --- |
| HIF-1alfa | 1/40 | NovusBio – NB100-105SS |
| HSP-90 | 1/400 | Santa Cruz – SC7947 |
| NOS2 | 1/500 | Santa Cruz – SC 651 |
| eIF-4E | 1/400 | Santa Cruz – SC 9976 |
| P eIF-4E | 1/50 | ThermoFisher – Ser209 |
| NFK-beta | 1/100 | Santa Cruz – SC 372 |

### Statistical Analysis

The Kolmogorov-Smirnov normality test with Dallal-Wilkinson-Lille for p values was used to evaluate the Gaussian distribution of data values. The t-test was used, and the values were expressed as mean ± standard deviation (SD). The Welch test was performed to correct situations of heteroscedasticity, that is when was observed a large variance between groups studied. p<0.05 was considered significant. GraphPad Prisma v. 01 software (GraphPad Software, Inc., USA) was used for statistical analysis and graph construction.

## RESULTS

### Body mass, renal area, and CM cells number

The male 17-GD LP offspring showed a significant reduction in the body mass compared to the age-matched NP group (Figure 1). The kidney/body mass was also reduced in 17-GD (Figure 1). In the 17-GD LP animals, a significant reduction (7.6%) in the renal area was observed. In addition, the calculated nephrogenic cortical area was 31% reduced (LP= 27.5 ± 1.2 vs. NP = 58.1 ± 1.6, N=4 of each, p<0.0001), and the medullar 34% enhanced (LP= 72.5 ± 1.3 vs. NP= 42 ± 1.6, N=4 of each, p<0.0001) compared to NP progeny. The CMs presented an increase in both areas (103%) and several Six2 positive cells (32%) in the 17GD LP kidneys. We did not observe differences between the (Figure 1) regarding placental mass. The areas occupied by CAP and comma-shaped vesicles were significantly smaller in offspring from the LP group compared to NP progeny (Figure 1).

**Figure 1.**
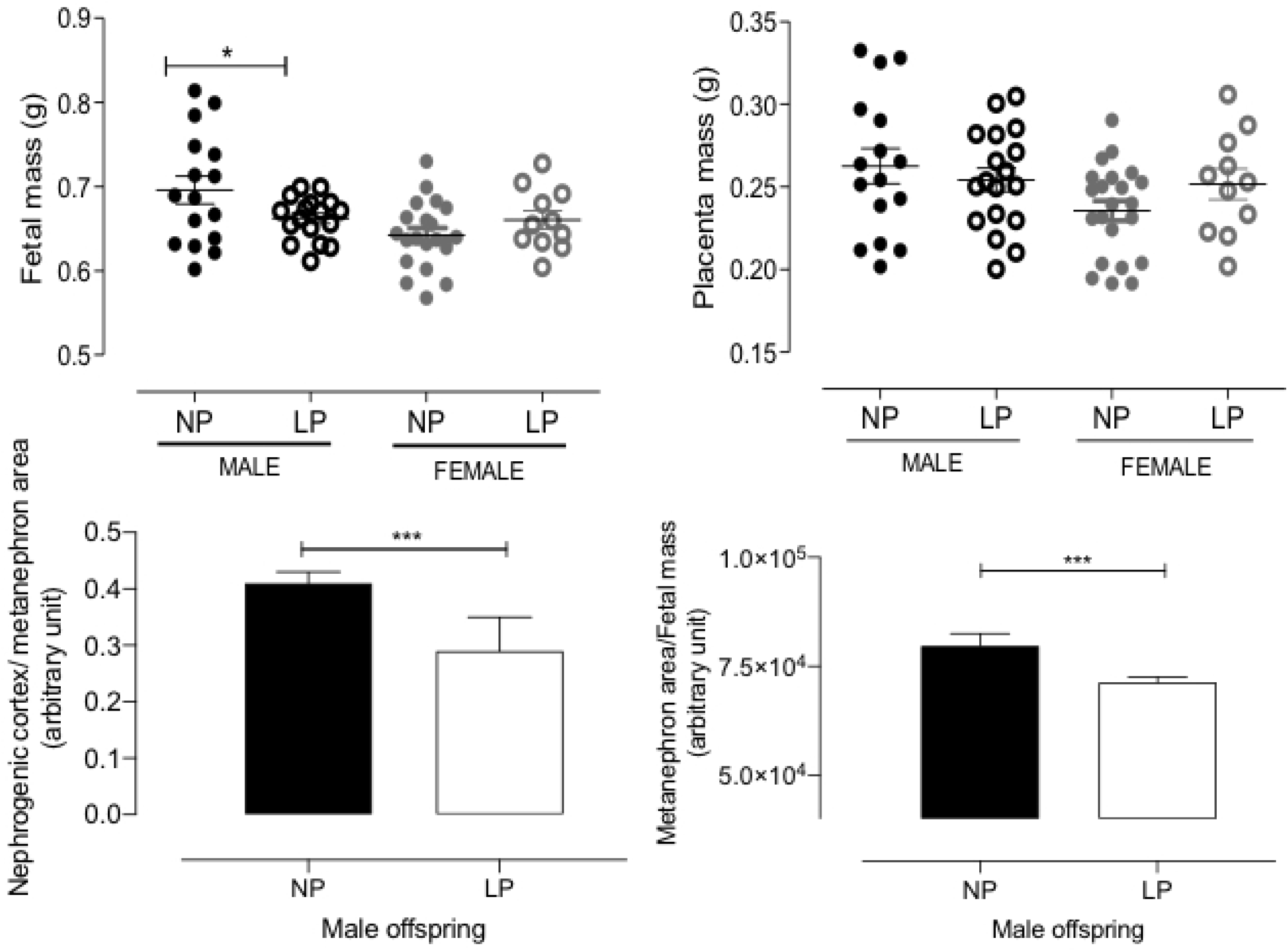
The graphics represent the female and male 17-DG fetal body, placenta masses, and nephrogenic cortex/metanephros area ratio in male NP progeny compared to LP offspring. Mean ± SD, *p<0.05, ***p<0.0001.

### Gene expression analysis

Figure 2 shows the mRNA expression of the elected predicted gene expression analysis in metanephros of males in 17-DG. We observed a significant increase in TGFβ, mTOR, elF4, HSP90, p53, p300, NFκβ, and AT2 compared to age-matched NP (Figure 2). Regarding HIF-1α, the LP animals showed an increase in the expression of their mRNA concerning the NP. However, we did not observe statistical significance (p=0.053). The presentation of mRNAs encoding NOS2, AT1⍰1, and AT1⍰1 although, AT2 receptors have been different between groups, was not altered in LP animals compared to control animals (Figure 2).

**Figure 2.**
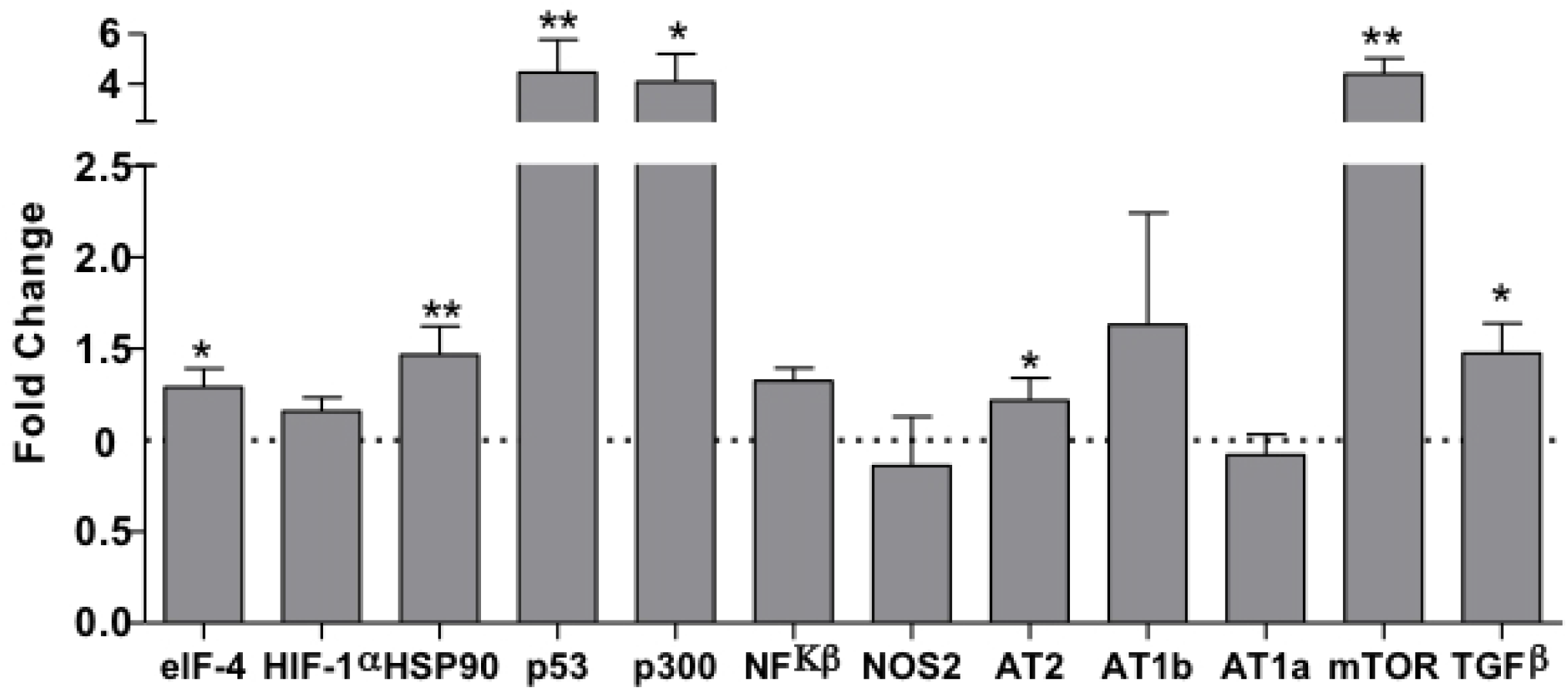
Renal expression estimated by SyBR green RT-qPCR of mRNA from a fetal kidney of the 17-GD LP offspring. The expression was normalized with GAPDH. The authors established a cutoff point variation of 1.3 (upwards) or 0.65 (downwards), and data are expressed as fold change (mean ± SD, n = 5) concerning the control group. * p≤0.05: statistical significance versus NP.

### Immunohistochemistry

As depicted in Figure 3, the mTOR is localized in tubular segments below the nephrogenic zone was increased in 17-DG LP compared to age-matched NP offspring. The intensity of TGFβ1 reactivity was significantly enhanced in all renal tissue in LP than in NP (Figure 4). The HIF-1α is widely shown in cortical and medullar areas of the metanephros (Figure 5). The staining presents higher labeling in 17-DG offspring, preferentially located in the nuclear site associated with low cytosolic intensity than age-matched NP offspring (t=0,9669, df=85, p=0.0001). elF4 immunoreactivity was dispersed in the cytosol and nuclei of metanephros cell types (Figure 6). Although no difference was observed between immunostained cells from NP and LP progeny, we observed a significant reduction in LP progeny kidney tissue (t=5.838, df=101, p=0.0001). Additionally, studying the phosphorylated form of elF4 staining was observed increased reactivity in the scattered mesenchymal cells (t=7,486, df=89, p=0.0001). However, the immunoreactivity area and the labeling intensity in the CAP revealed a significant reduction in LP compared to NP (Figure 7). In metanephros of NP animals, the HSP90 protein was located in small intensity in the cytosol of different cell types. Otherwise, endothelial cells are easily showed higher immunostaining to HSP90 (Figure 8). The immunoreactivity for HSP90 is higher in different metanephros nuclear cell types in LP offspring (t=6,770, df=115, p=0.0001). Through quantification, increased intensity and percentage of marked CAP area in LP progeny were observed compared to the NP offspring (Figure 8). In control animals (NP), the NFκβ staining is weak in all metanephric tissue (Figure 9). However, in 17-DG LP, the NFκβ and nuclear immunoreactivity are significantly enhanced in all cells (t=2,822, df=117, p=0.0056). Also, the percentage of labeled CAP area was increased considerably in 17-DG LP (Figure 9), and the immunostained intensity, although higher in micrograph 9, was not different in both CAP groups (Figure 9). Immunoperoxidase staining for NOS2 was not observed in metanephros from NP animals (Figure 10). Although significantly increased in LP progeny metanephros, the NOS2 immunostaining occurred weakly throughout all metanephros extent (t=4,482, df=126, p=0.009). Through the quantifications, we obtained a considerable increase in the percentage of marked CAP area and in the intensity of marking in the CAP (Figure 10) in LP offspring compared to NP progeny.

**Figure 3.**
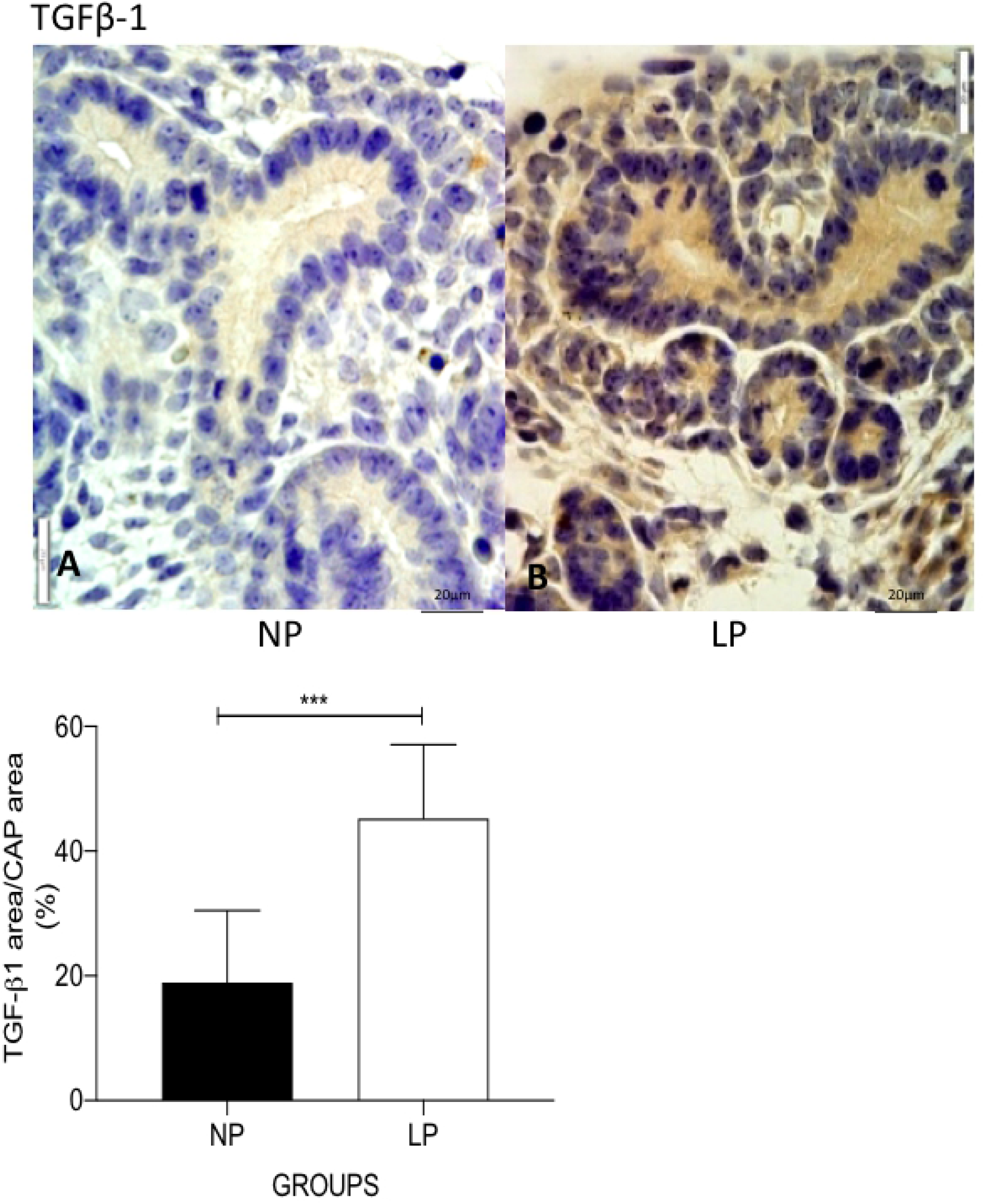
TGFβ-1 immunoperoxidase in kidneys of 17-GD LP progeny compared to NP offspring. The graphics represent the TGFβ-1 immunostained area/CAP area ratio in male 17-DG LP compared to age-matched NP progeny. Mean ± SD, ***p<0.0001.

**Figure 4.**
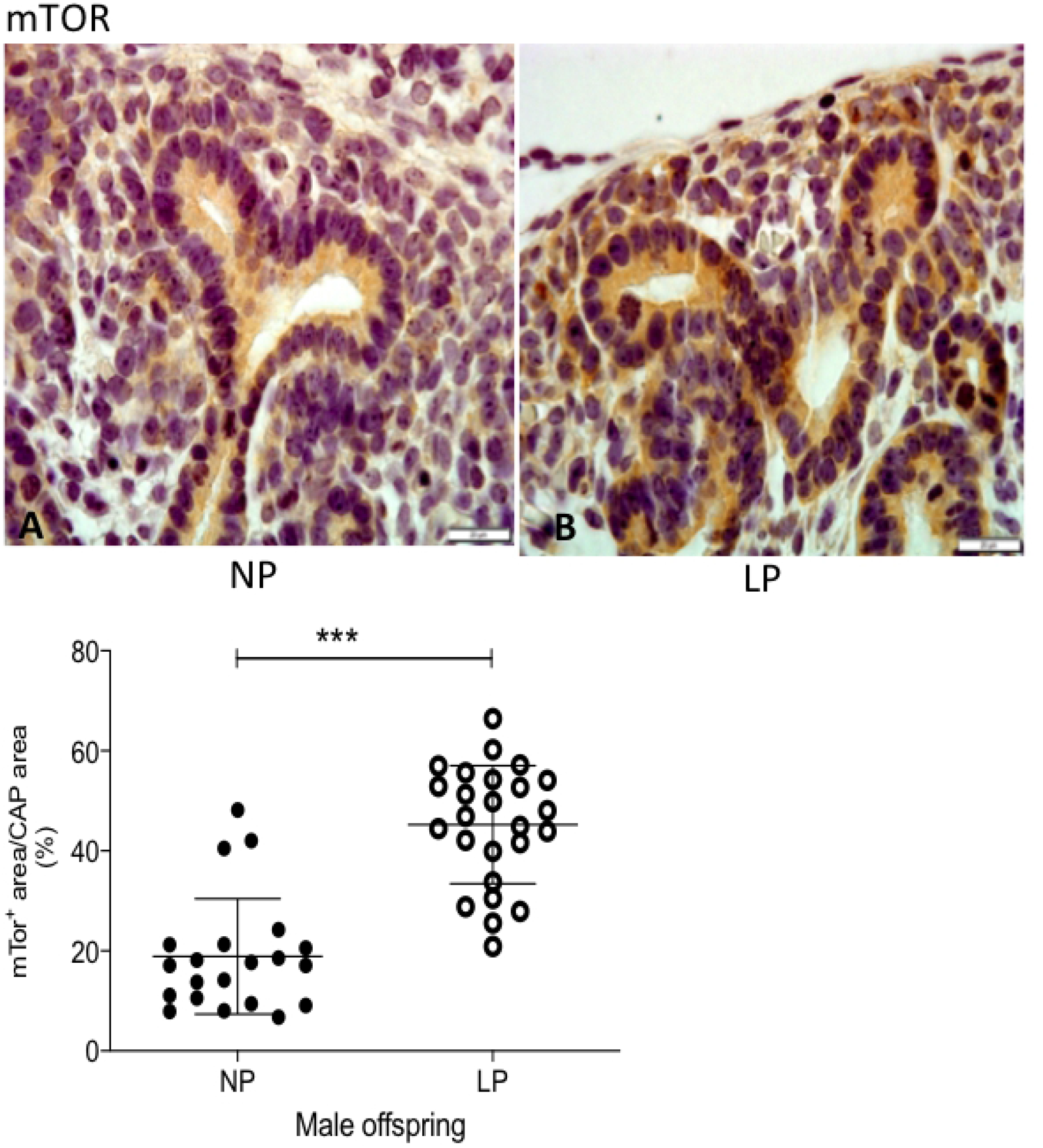
mTOR immunoperoxidase in kidneys of 17-GD LP progeny compared to NP offspring. The graphics represent the mTOR immunostained area/CAP area ratio in male 17-DG LP compared to age-matched NP progeny. Mean ± SD, ***p<0.0001.

**Figure 5.**
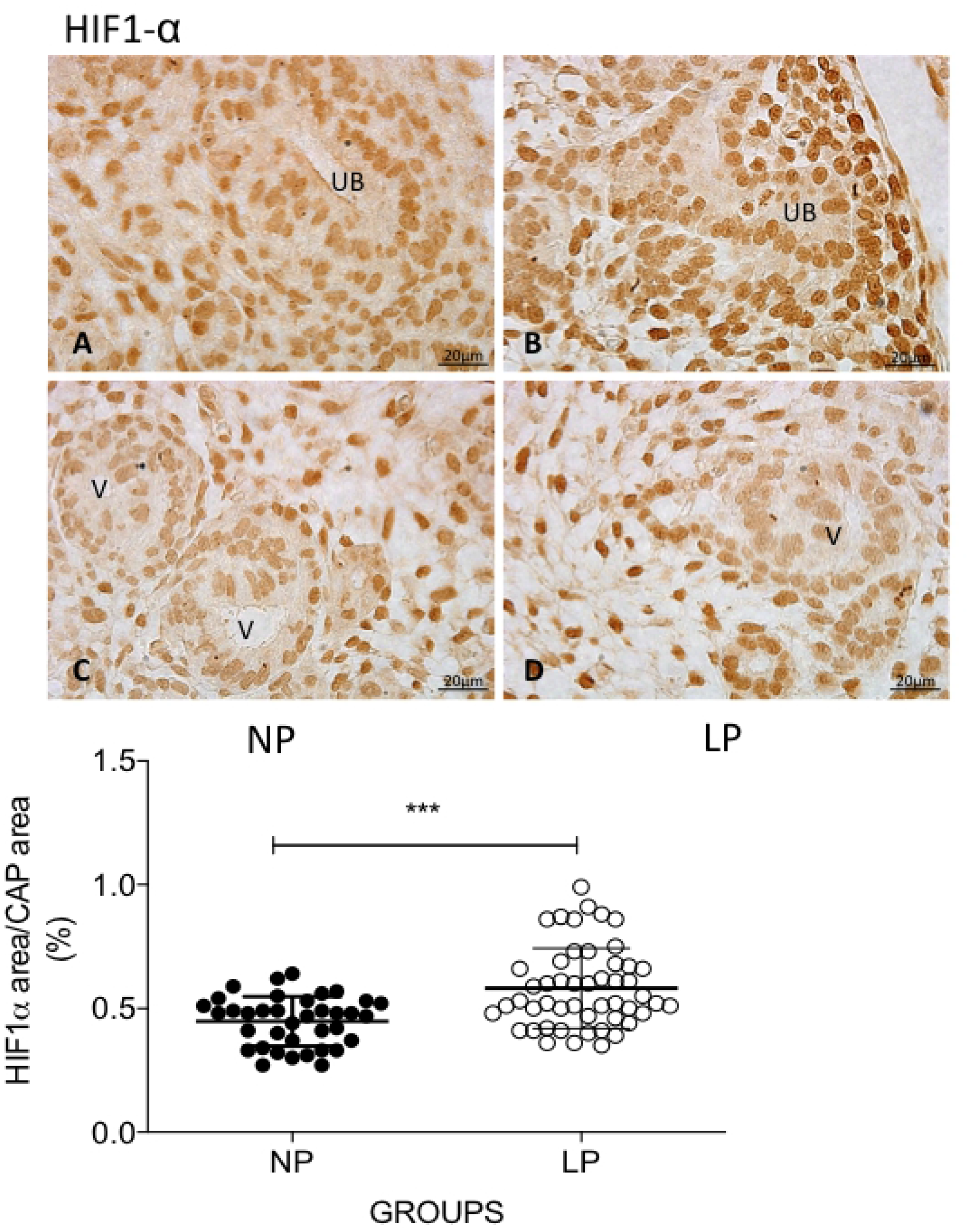
HIF-1α immunoperoxidase in kidneys of 17-GD LP progeny compared to NP offspring. The graphics represent the HIF-1α immunostained area/CAP area ratio in male 17-DG LP compared to age-matched NP progeny. Mean ± SD, ***p<0.0001.

**Figure 6.**
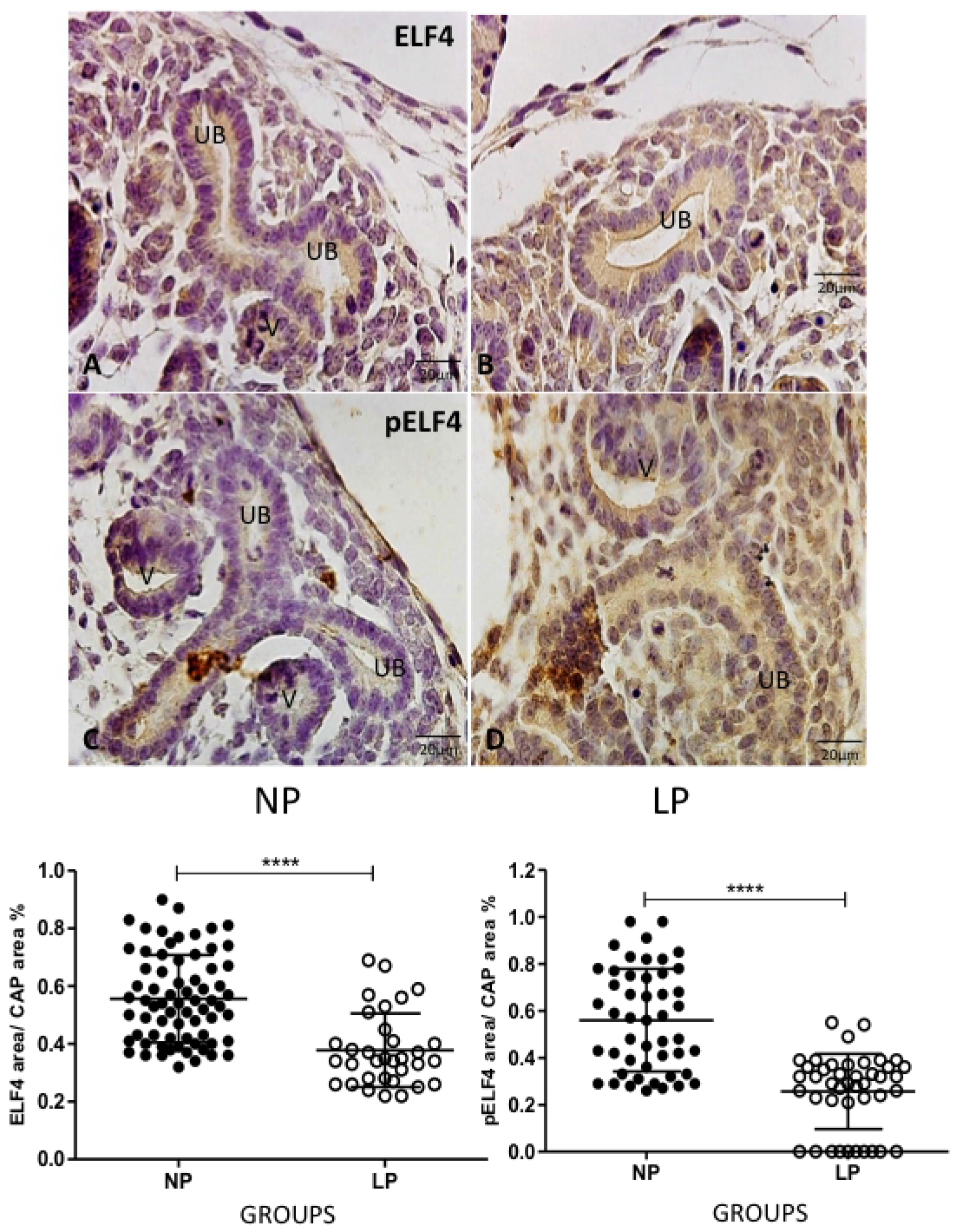
ELF4 and pELF4 immunoperoxidase in kidneys of 17-GD LP progeny compared to NP offspring. The graphics represent the ELF and pELF immunostained area/CAP area ratio in male 17-DG LP compared to age-matched NP progeny. Mean ± SD, ****p<0.0001.

**Figure 7.**
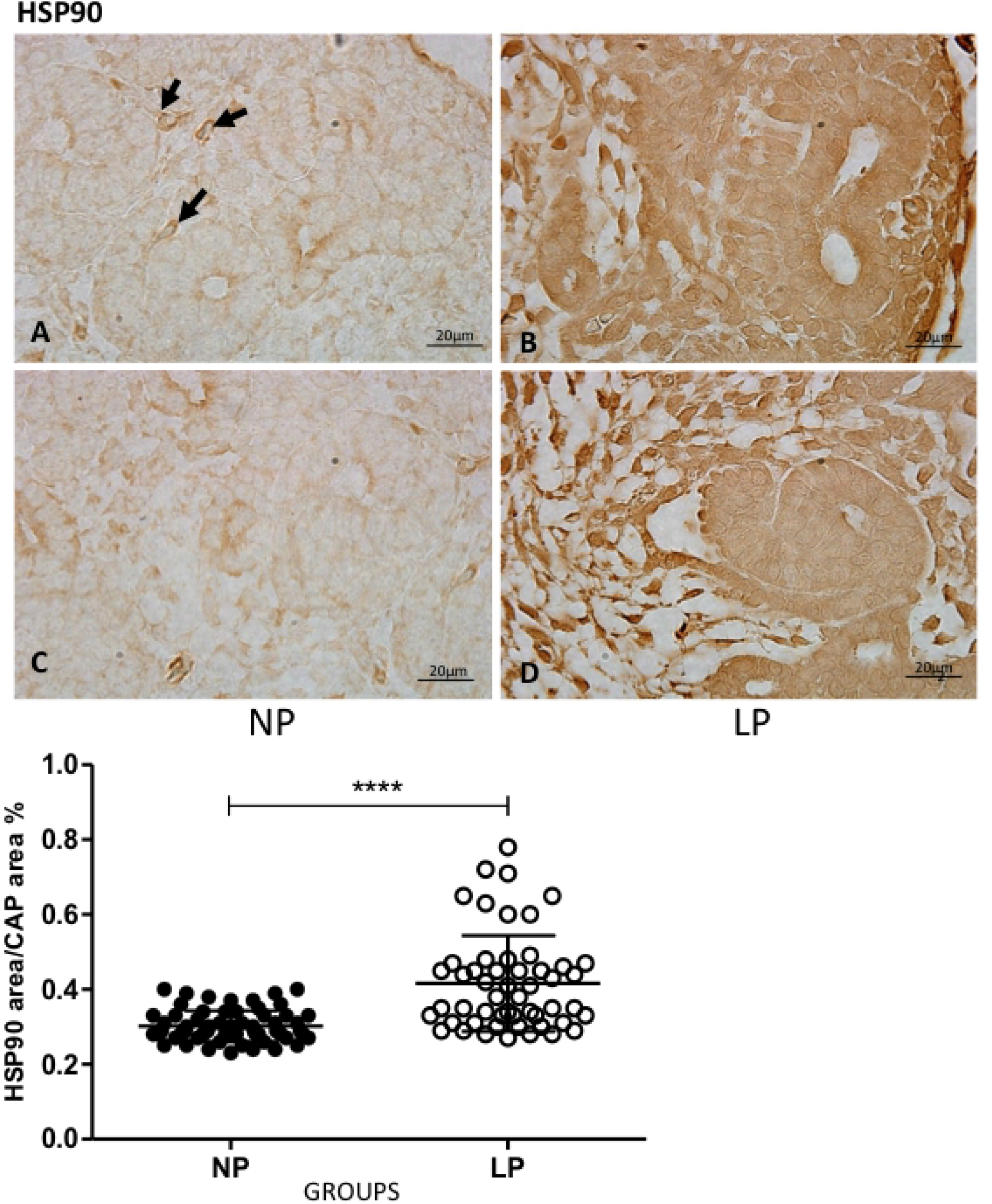
HSP90 immunoperoxidase in kidneys of 17-GD LP progeny compared to NP offspring. The graphics represent the HSP90 immunostained area/CAP area ratio in male 17-DG LP compared to age-matched NP progeny. Mean ± SD, ****p<0.0001.

**Figure 8.**
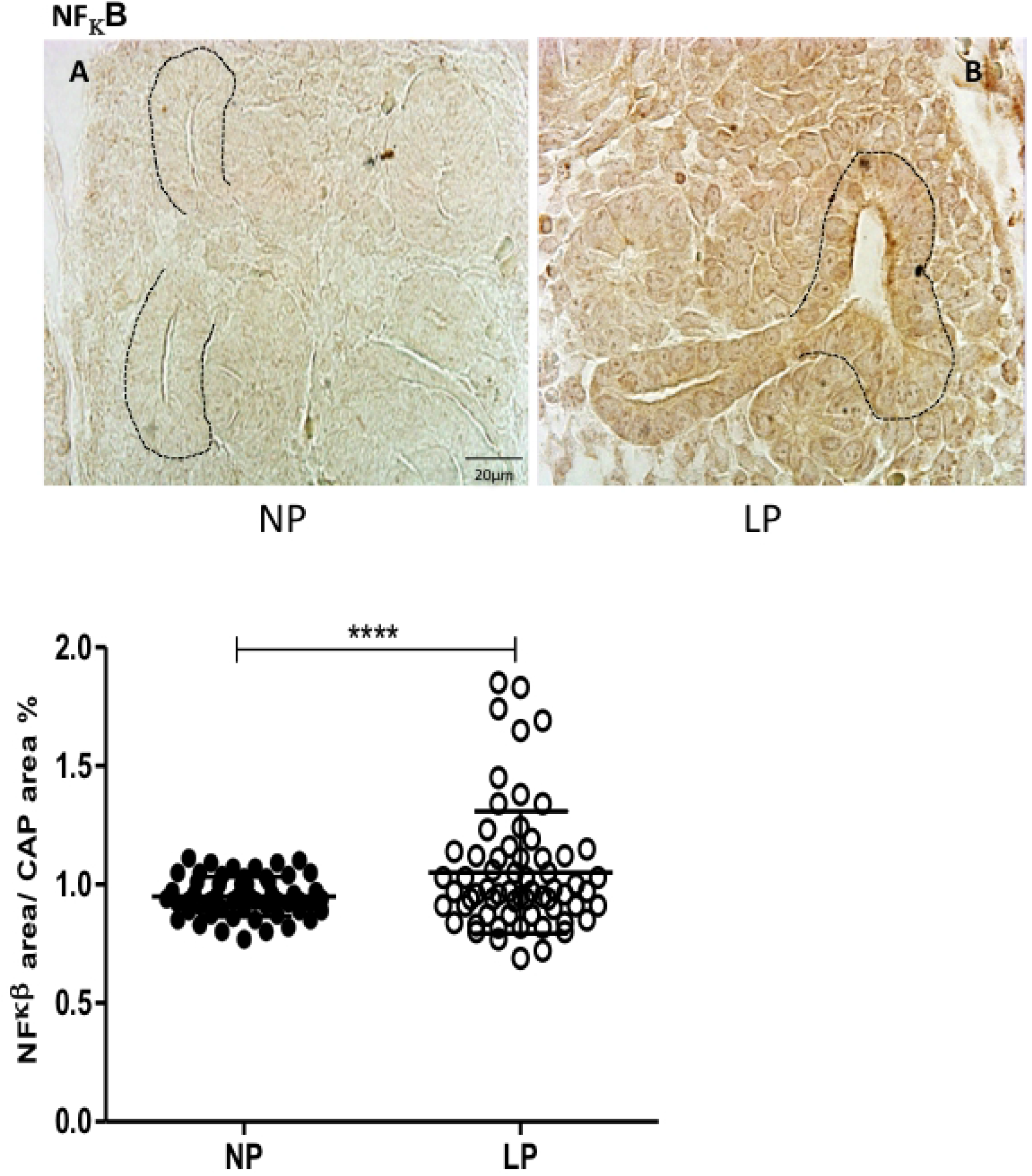
NFκβ immunoperoxidase in kidneys of 17-GD LP progeny compared to NP offspring. The graphics represent the NFκβ immunostained area/CAP area ratio in male 17-DG LP compared to age-matched NP progeny. Mean ± SD, ****p<0.0001.

**Figure 9.**
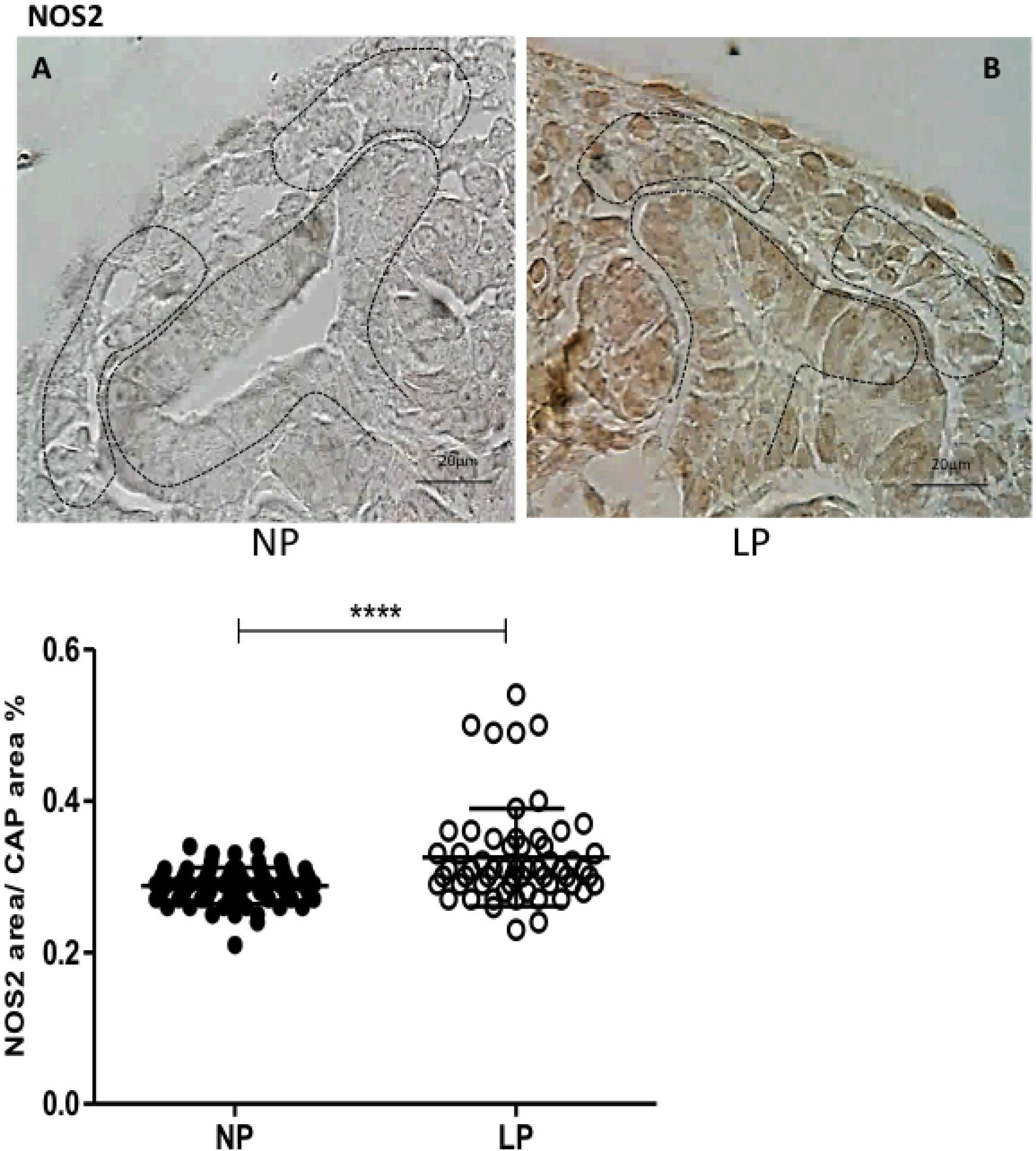
NOS2 immunoperoxidase in kidneys of 17-GD LP progeny compared to NP offspring. The graphics represent the NOS2 immunostained area/CAP area ratio in male 17-DG LP compared to age-matched NP progeny. Mean ± SD, ****p<0.0001.

**Figure 10.**
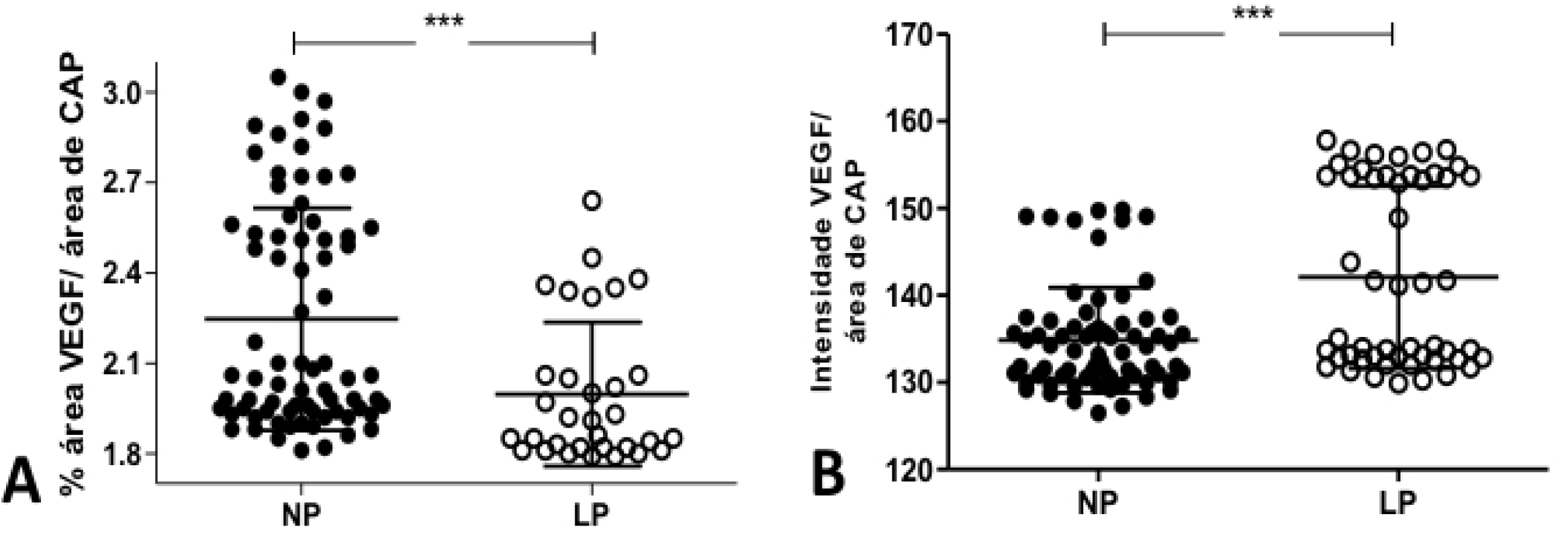
The graphics represent the VEGF immunostained area/CAP area ratio in male 17-DG LP compared to age-matched NP progeny. Mean ± SD, ****p<0.0001.

## DISCUSSION

Assuming that chronic diseases in adulthood are “programmed” during the embryo-fetal period, the study of triggering factors for this programming is of great importance. Nephrogenesis depends on adequate progenitor cells’ self-renewal, survival, proliferation, and differentiation capacity during perinatal kidney development in rats. Exist a consensus that miRNAs drive renal growth-regulating gene expression of proteins involved in key signaling pathways [5,7,36–38]. Recently we established miRNA and elected target mRNA expression in maternal low-protein intake 17-GD male metanephros [5,7]. The current study depicted the contribution of previous studied and elected miRNAs transcripts during nephrogenesis in the maternal low protein intake model by data from this age in 17-GD fetus kidneys from LP progeny compared to NP offspring [7].

Studies in our laboratory reveal that gestational caloric and protein restriction led to the reduced number of nephrons observed during the fetal period and confirmed in adult life, probably secondary to a decrease in the number of progenitor cells [3,4,8,39]. The cellular mechanisms responsible for the impairment of renal outcome in programmed animals led to evidence of a complex network of protein and miRNA interactions involved in renal organogenesis. Therefore, the evaluation of the HIF-1α signaling pathway was based on the prior observation of altered proteins of this pathway in metanephros of programmed animals in our Lab, using the same experimental design of the present work [5–7]. This observation aims to elucidate particular HIF mRNA transcripts and possibly the relationship of this transcript in the reduced nephron number observed in this severe protein-restricted model [3,4].

Preliminary data showed that a low-protein diet during embryonic development increased levels of TGFβ1/2 in metanephros of 17-DG fetuses [7]. Also, it demonstrated a significant increase in mTOR in the maternal restricted protein progeny [7]. It has been shown that elevation of mTOR expression promotes activation via PI-3K/AKT pathway inducing the expression of HSP90 and, consequently, contributing to the stabilization and translocation of HIF-1α to the cell nucleus. In a previous study [7], among miRs, the miR-199a-5p expression was the most significantly related to mTOR mRNA encoding and protein immunoreactivity. That mTOR mRNA expression has a pronounced reduction in the 17-GD LP offspring kidney.

Additionally, the upstream of mTOR mRNA transcript and TGFβ-1 protein reactivity were upregulated at the 17-GD LP compared to age-matched NP progeny. mTOR signaling plays a central role in sensing response to intracellular nutrient availability [40]. Another proposed mechanism for the increase in HIF-1α involves the PI-3k/AKT signaling pathway, which would be its activation by TGFβ, leading to mTOR enhancement, as previously mentioned. The kidney’s transcriptome from a fetal baboon, whose mothers are submitted to nutrient-restricted intake corroborated with a demonstration of the mTOR signaling pathway be central to a reduction in the nephron number in this model [40]. Although it is widely known that mTORC1 has an essential role in embryo development, keep completely unclear the complex mechanisms in stress conditions [41]. Hudson et al., 2002 [42] defined a relationship between the increased mTOR pathway activity and HIF-1α, who treated cell cultures with an mTOR inhibitor.

For decades, as mentioned above, the relationship between HIF-1α and HSP90 was observed in vitro. It has been demonstrated that a fall in HSP90 expression decreases in HIF-1α tissue levels, promoting a reduction in its transposition to the cell nucleus and binding in target genes [43–47]. On the other hand, the low concentration of HSP90 also promotes an increase in VHL protein, leading to ubiquitination and degradation of HIF-1α in proteasomes and consequent inhibition of its transcriptional activity. However, a non-hydroxylation of HIF-1α prevents its binding to HSP90 avoids the destruction of HIF-1α by unspecific degradation pathways. The present study demonstrated the high levels of HSP90 in 17-GD LP progeny kidneys that prevent the degradation of HIF-1α, accelerating its cellular accumulation and thus allowing its transcriptional activity [47].

Several proteins have already been described as participating upstream and downstream pathways of HIF-1α. Among these, NFκβ has been identified. When activated, NFκβ translocated to the cell nucleus, where they bind to DNA and regulate the transcription of their target genes. Studies have demonstrated that components of the HIF-1α and NFκβ signaling pathways share some of their genetic targets. van Uden et al. (2008), showed that stimulation of NFκβ by TNFα also led to an increased level of HIF-1α mRNA. Under different circumstances, these findings confirm the results of the current study in 17-DG LP compared to NP fetal kidney tissues, since the gestational low-protein diet also promoted a significant increase in mRNA expression and NFκβ protein levels and, consequently, in HIF-1α [48].

The p53, a protein with a molecular weight of 53 kDa, is encoded by chromosome 17 as a tumor suppressor protein. It plays a primary and essential role in maintaining the integrity of the genetic code. The p53 keeps the same sequence of nucleotides throughout the entire DNA molecule by repairing proteins to restore the DNA to its normal state or to induce the synthesis of repair proteins maintaining DNA structure constant in all organism cells. Several studies have demonstrated that p53 down-regulated HIF-1α through evidence that hypoxia-induced stress can increase p53, consequently promoting a decrease in transcriptional HIF-1α. The relationship between the tumor suppressor p53 and HIF-1α is still unclear. Both HIF-1α and p53 need to bind Ep300 for their respective transcriptional activities. Therefore, evidence shows that, when there is severe hypoxia, there is an increase in p53, which would lead to an increase in the degradation of the HIF-1α protein [32,33,49–51]. Likewise, the reduction of p53 would cause an increase in the stability of HIF-1α and an increase in its transcriptional activity.

On the other hand, some authors postulated that the increase in p53 in hypoxia would not be associated with low O2 tensions, per se, but with the change in the environment caused by hypoxia – such as decreased pH and lower glucose availability. These changes would lead to increased p53 and could be related to the adverse fetal development environment [32,50,52,53]. The cellular concentration of p53 must be tightly regulated. Thus, the high level of p53 may suggest an acceleration of the aging process by excessive apoptosis of metanephric mesenchymal progenitor cells. However, the present study showed an increased p53 mRNA associated with an increase in HIF-1α, suggesting an opposite relationship with the previously described negative regulation of HIF-1α by p53. This evidence leads us to conclude that the relationship between p53 and HIF-1α needs to be further clarified. Ep300 is a coactivating molecule, central to the protein-protein interaction network that links HIF-1α to other signaling pathways, which allows the specific response of HIF-1α and its transposition to the nuclei of cells and the regulation of their multiple target genes. The basis of the link between Ep300 and HIF-1α is the inhibition of hydroxylation of HIF-1α. In environments with normal oxygen tension, HIF-1α is hydroxylated, which prevents its binding to Ep300 and, consequently, its transcriptional activity. In the current study, an increase in Ep300 expression was seen in agreement with the rise in HIF-1α, suggesting that in an adverse environment, caused by gestational protein restriction, interfering with renal development causes inhibition of hydroxylation and an increase in the binding between HIF-1α and Ep300, and transposition to the nucleus [54–58].

Signaling pathways modulated by growth factors, such as PI-3k/AKT/PKB and MAPK pathways, stabilizing and activating HIF-1α expression in a typical environment. On these circumstances, HIF-1α is activated by several growth factors and cytokines, including IGF-1 and IGF-2, insulin, interleukin-1β, follicle-stimulating hormone, Ang II, TGFβ, and TNFα. Several components of the PI-3/AKT signaling pathway have also been shown to activate HIF-1α. In the same sense, cells with regular AKT activity present HIF-1α induction in an environment with usual oxygen tensions, while the treatment of cells with PI-3 inhibitors blocks the expression of HIF-1α. [59–61]. Conversely, some authors argue that PI-3k/AKT signaling is not involved in the induction of HIF-1α in inadequate O2 tissue tension. Thus, the function of this activation has not yet been precisely established [62–64].

It is believed that the mechanism of stimulation of HIF by growth factors occurs predominantly at the level of protein synthesis than by mRNA transcription. It showed that HIF-1α mRNA remained unaffected by IGF1 stimulation, which keeps HIF-1α ubiquitinated and capable of binding to VHL protein. Also, it has been revealed those levels of HIF-1α protein, the HIF-DNA sequence binding, and its transcriptional activity were secondary to the protein synthesis of HIF-1α and not through the inhibition of HIF-1α its degradation. In parallel, the accumulation of HIF-1α by stimulating TNF-α and NFκβ, as seen above, appears to be mediated by protein synthesis of HIF-1α [60,61,64,65].

Ang II acts on cell activity differently, either through stimulating cell growth, hypertrophy, or apoptosis, depending on the specific situation. The binding of angiotensin II with its AT1 receptor leads to the activation of intracellular second messengers, consequently increasing nitric oxide production through phosphorylation caused by NOS-2, which in a counter-regulatory manner inhibits the cellular hypertrophy caused by Ang II. Knockout mice for HIF-1α showed increased expression of the AT1 receptor in the vasculature and, consequently, increased angiotensin effects, including increased blood pressure [66,67]. As mentioned above, the HIF-1α activation pathway coincides with the phosphorylation of p53, indicating that HIF-1α phosphorylation is, at least in part, secondary to the upregulation mechanism by NO [68–72]. These authors demonstrated that variations in NO concentration would determine a positive or negative effect on HIF-1α. They described that in environments with elevated concentrations of NOS and, consequently, of NO, the stabilization of HIF-1α occurs. However, in a low concentration of NO, there is a significant decrease in the HIF-1α levels [73 75]. It is believed that degradation of HIF-1α would be secondary to the inhibition of mitochondrial respiration, which leads to a local increase in oxygen and activate the prolyl hydroxylase (PHD) enzyme. This mitochondrial function would be responsible for forming reactive oxygen species (ROS), triggering events that would start HIF-1α and promote cellular adaptation and stabilize and increase the activity of HIF-1α [73–77]. Thus, the relationship between activated HIF-1α by NO and, consequently, by NOS-2 is evident in the literature. The present study observed an increased NOS-2 expression in the kidney of 17-GD low-protein diet offspring. Based on these findings, we could suggest that the intrauterine development environment exposed to a low-protein diet would stabilize HIF-1α, promoting their migration to the cell nucleus.

One of the target genes of HIF-1α, VEGF, is necessary for vascular, tubular, and glomerular cell hypertrophy and proliferation and can be activated by inflammatory and pro-fibrotic factors, including Ang II. It is believed that the cellular effects of Ang II may be through the activation of HIF-1α and its growth factor target genes [78]. The same experimental model has shown that miR 199a-5p is related to activation of the WNT pathway regulating vascular and nephron development [7]. Here, we may suppose that the previous finding demonstrated that miR-199a-5p positively modulated VEGF expression (mRNA and protein) in the fetal LP kidneys. VEGF signaling is a downstream event of the mTOR pathway, but, in the present study, the VEGF mRNA was not accompanied by the mTOR mRNA encoding; however, it occurs parallel to VEGF immunoreaction. Kitamoto and col (1997) studied in vivo the role of VEGF in kidney development by blocking the endogenous VEGF activity in newborn mice [79]. They showed a reduced nephron number and abnormal glomeruli. The increased VEGF reactivity observed in 17-GD kidneys from the LP progeny indicates a possible compensation of peritubular and glomerular capillary development; once in the 17-GD LP kidneys, the low expression could impair vascular development.

In addition, in studies with tubular epithelial cells, Ang II, through the increase of HIF-1α and promoted an increase in VEGF expression by reducing prolyl-hydroxylase 3 (PHD) enzyme concentration that promotes a decreased HIF-1α degradation. Likewise, Ang II infusion in mice led to overexpression of VEGF and increased transcriptional activity of HIF-1α in the kidney [80–86]. The reduced HIF-1α in fetal renal tissue causes a decreased VEGF expression [87] and is undoubtedly one factor that reduces the number of renal functional units. In the current study, the enhanced HIF-1α immunostained may increase AT2 receptors and VEGF mRNA expression, confirming studies mentioned above, which could secondarily be responsible for the increase of expression of HIF-1α and its molecular components of its pathway.

In studies from our laboratory in the same pattern of fetal programming, which would, in turn, was demonstrated inhibition of the transcriptional factor elF4, as has been shown previously consequently, to activation of HIF-1α [5,7]. This inhibition reduced the stabilization of HIF-1α and its transcription, regardless of the environment with low or normal oxygen tension. The evidence indicates that mTOR may act simultaneously as a positive stimulator of HIF-1α and an inhibitor of elF4 expression. This finding is corroborated by a previous study, which showed an increase in mTOR in animals submitted to fetal programming with a low-protein diet. The current study results align with this evidence, showing and proving an elevation of HIF-1α. [5–7,42].

In conclusion, the current study data supported that nephron onset impairment in 17-DG fetuses kidney, programmed by gestational low-protein intake is, at least in part, related to alterations in the HIF-1α signaling pathway. Factors that facilitate the transposition of high levels of HIF-1α to the mesenchymal cell’s nucleus, such as increased expressions of NOS, Ep300, and HSP90, may have an essential role in this regulatory system. This alteration leads to the inhibition of adaptive responses to the adverse environment, secondary to an increase in ungraded HIF-1α, possibly associated with a reduction in the transcription factor elF-4 and proteins of their respective signaling pathways. Although surprising, the high expression of p53 may be necessary for the apoptotic process that possibly occurs, culminating in the reduction of proliferation and early elevation of cell differentiation in this experimental model (Figure 11). Consequently, we may suggest an early maturation process of renal cells, inhibition of nephron progenitor cell division, and reduction of renal functional units in the offspring of rats submitted to severe gestational protein restriction.

**Figure 11.**
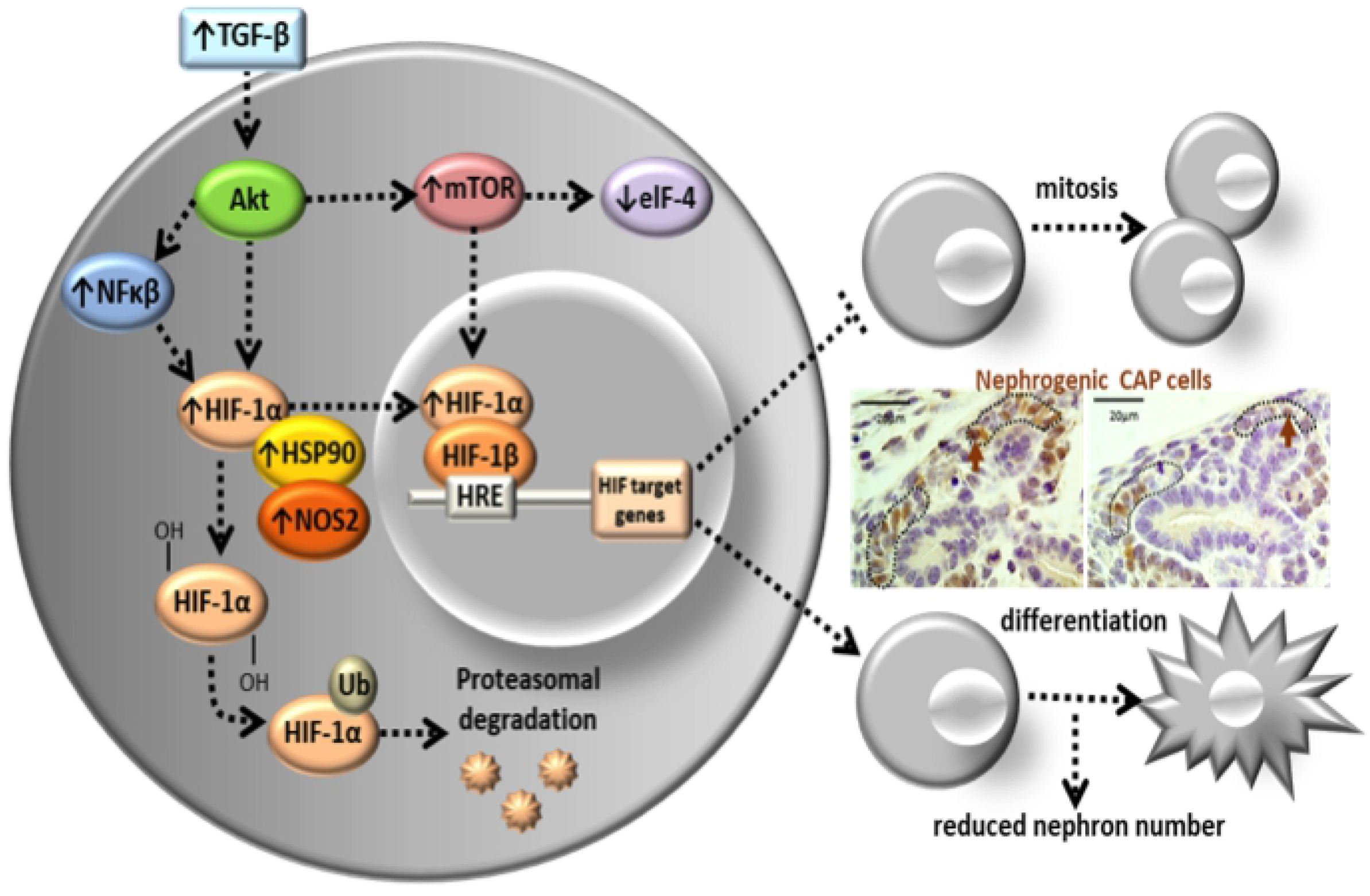
Schematic diagram of fetal kidney development in gestational protein-restricted offspring. The figure depicted the nephron onset impairment in 17-DG fetuses kidney, programmed by gestational low-protein intake, related to canonical pathways alterations in the HIF-1α signaling pathway. Expressions of NOS, Ep300 potentiate the transposition of HIF-1α to the mesenchymal cell’s nucleus, and HSP90, possibly associated with a reduction in the transcription factor elF-4 and proteins their respective signaling pathways. These results may suggest an early maturation process of renal cells, inhibition of nephron progenitor cell division, and reduction of renal functional units in the offspring of rats submitted to severe gestational protein restriction.

## LIST OF ABBREVIATIONS

Ang II: Angiotensin II
AT1 and AT2 receptors: Type 1 and Type 2 angiotensin receptor
BSA: Bovine serum albumin
cDNA: complementary deoxyribonucleic acid
CEUA/UNESP: Institutional Ethics Committee
CM: metanephros cap
DAB: 3,3’-diaminobenzidine tetrahydrochloride
DNA: deoxyribonucleic acid
elF4: E74-like factor
pelF4: phosphorylated E74-like factor
EP300: E1A binding protein p300
PCR: Polymerase chain reaction
GD: gestational days
GAPDH: Glyceraldehyde 3-phosphate dehydrogenase
HIF: Hypoxia-inducible factor
HSP90: heat shock protein 90
IDT: Integrated DNA Technologies
IGF: Insulin-like growth factor
LP: gestational low-protein intake
CM: Metanephros cap
MM: Metanephros mesenchyme
Map2k2: mitogen-activated protein kinase kinase 2
*miRNA (miR)*: a small non-coding RNA molecule
*miRNA-Seq*: *miRNA transcriptome sequencing*
mRNA: messenger ribonucleic acid
mTOR: mammalian target of rapamycin
NFκβ: nuclear factor kappa-light-chain-enhancer
NGS: Next Generation Sequencing
NO: Nitric oxide
NOS: nitric oxide synthetase
NOTCH1: single-pass transmembrane receptor protein
NP: normal protein intake
PHD: Prolyl-hydroxylase 3 enzyme
RIN: RNA Integrity Number
RT-qPCR: reverse transcription-polymerase chain reaction quantitative real-time
TGFα and TGFβ: α and Beta transformer growth factor
TGFβ-1: transforming growth factor-beta 1
TNF: Tumor necrosis factor
UB: ureter bud
U6 and U87: internal reference gene
VEGF: endothelial vascular growth factor
VHL E3: tumor suppressor Von Hippel Lindau
17-GD: 17th gestational day

---

**Table S1 (Supplemental Information).** *The miRNA sequencing data*

